# MiR-17-5p regulates proliferation and apoptosis of uterine fibroids via targeting ESR1

**DOI:** 10.1101/2020.11.10.376384

**Authors:** Lan Luo, Jian-Jun Cao, Hui-Min Zhang, Kun Chen, Xing-Hua Liao, Kun Li

## Abstract

The treatment of uterine fibroids and the development of new drugs depend on adeeper understanding of the developmental mechanisms of uterine fibroids. Here, the role of ESR1 and miR-17 on the uterine fibroids cell proliferation and apoptosis and their relationship were investigated in USMCs. Our results showed that ESR1 increased the proliferation of USMCs and inhibited their apoptosis. In addition, ESR1 could directly bind the promoter regionof TP53 and inhibit its expression. MiR-17 increased the apoptosis of USMCs and inhibited their proliferation via decreasing the level of ESR1 by targeting its 3’UTR. Our research provides a new understanding of the development of uterine fibroids and provides a theoretical basis for the treatment of uterine fibroids.

## Introduction

Uterine fibroids are benign lesions or uterine tumors.[1] At present, the main treatment of uterine fibroids is hysterectomy, which may affect fertility and have a profound impact on health.[2] The development of new uterine fibroids treatments and drugs depends on a deeper understanding of the development of uterine fibroids and the discovery of new therapeutic targets.

Hormone factors synthesized by estrogen are indispensable in the progression of cancer. ESR1 is a transcription factor that responds to estrogen and various growth factors in a variety of tissue types and is activated by ligands, further affecting downstream gene expression, including affecting the mammary gland Proliferation of epithelial tissue.[3] The function of ESR1 in uterine fibroids has not been reported.

MiRNAs can mediate mRNA translational repression or cleavage and down-regulate the protein levels by binding to the complementary site in 3’UTR or mRNA.[4–6] MiR-93 and miR-17 belong to the miR-17-92 family, which share similar binding sites. Previous studies have shown that miR-93-5p exerts tumor suppressive effects in breast and cervical cancer.[7–10] Whether miR-17 can inhibit the occurrence and development of uterine fibroids is still unclear.

The roles of ESR1 and miR-17 on the proliferation and apoptosis were detected in USMCs in this study. Our data showed that ESR1 could promote proliferation and inhibit apoptosis of USMCs, ESR1 directly bind the promoter of TP53 to inhibit its expression. MiR-17 increased apoptosis of USMCs and inhibited their proliferation via inhibiting expression of ESR1 by targeting its 3’UTR. Our research provides a new understanding of the development of uterine fibroids and provides a theoretical basis for the treatment of uterine fibroids.

## Materials and Methods

### Cell Culture

USMCs were purchased from ATCC (USA). USMCs were cultured in DMEM (Invitrogen, USA) added with 10% FBS (Every Green, China) and incubate in a 37 °C incubator with 5% CO_2_.

### Plasmid constructs

To knockdown the endogenous expression of ESR1 (NM_001122741), three siRNAs: Si-ESR1-1:

5’-GACTATGCTTCAGGCTACCATTATG-3’, Si-ESR1-2:

5’-CGCTACTGTGCAGTGTGCAATGACT-3’, Si-ESR1-3:

5’-GACCGAAGAGGAGGGAGAATGTTGA-3’ were synthesized (Ribobio, China). And random sequence of the same length synthesized as a NC (Si-NC). The expression vector ESR1 could ectopically express ESR1, and the ESR1 cDNA fragment was inserted into pCDNA3.1 (Invitrogen, USA). The primer sequence for constructing the recombinant plasmid were as follows: ESR1 forward: 5’-GCGGGTACCATGACCATGACCCTCCACACCAAAGC-3’, reverse: 5’-GGCCTCGAGTCAGACCGTGGCAGGGAAACCCT-3’. For ectopic expression or knock down miR-17-5p (MIMAT0000070), miR-17-5p mimics and miR-17-5p inhibitor were synthesized (Ribobio, China) as well as the corresponding NCS, NC-mimics and NC-inhibitor. The ESR1 3’UTR WT fragment was cloned into dual-luciferase miRNA target expression vector pmirGLO (Invitrogen, USA), the primers used were as follows: ESR1-3’UTR-WT-luc forward: 5’-GCAGTCTAGAGGGCGCCAGGCAGGCGGGCGCCACCGCC-3’, reverse: 5’-AGGCTGGGGTTTTACCAGTTTTATTTCTAGACT-3’. The primers used for MUT ESR1 3’UTR (binding site of miR-17-5p was mutated) were as follows: ESR1-3’UTR-M-luc forward: 5’-GGGTGCAAGGAAAATTAGGGTACTCGCAAGAAGTTCGGTTCCGATGAATTCTTATCCC-3’, reverse: 5’-GGGATAAGAATTCATCGGAACCGAACTTCTTGCGAGTACCCTAATTTTCCTTGCACCC-3’. The promoter region of TP53 fragment were cloned into the luciferase vector pGL3-Promoter (Addgene, USA) (TP53-WT-luc). The primers sequence was as follows: TP53-WT-luc, forward: 5’-CGAGCTCTTACGCGTGCTAGCGGAAACACTTGGACTACAC-3’, reverse: 5’-AGATCTCGAGCCCGGGCTAGCTAGCGCCAGTCTTGAGCACA TGGG-3’. The primers used for MUT TP53 promoter region (binding site of ESR1 was mutated) (TP53-M-luc) were as follows: TP53-M-luc, forward: 5’-GCACTATTATCGGAGATGCAGGAACCGTCAGGAGCCCTA-3’, reverse: 5’-CTGCATCTCCGATAATAGTGCTGATAACATCTTGCGAATTTA TAA-3’.

### Transfection

All the mimics, inhibitor and the plasmids were transfected with lipofectamine 3000 (Invitrogen, USA). Briefly, the p3000 fraction and plasmid were incubated in a certain ratio, then incubated with lip3000 fraction, and the mixture was added to the cells. For mimics and inhibitors, incubate directly with lip3000 fraction and add to the guidance of the manufacturer’s instructions.

### Cell viability assay

We used the CCK-8 (Beyotime, China) to measure the cell viability. Concisely, 96-well plate was seeded with 2 * 10^4^ UMSCs per well. CCK-8 reagent was added to each well 48 hours after transfection, and detected the absorbance value at 450 nm using a full-wavelength microplate reader to calculate cell viability under the guidance of the manufacturer’s instructions.

### EdU-labeling assay

We used the Cell-Light TM EdU in vitro cell proliferation imaging kit (Ribobio, China) to detect the cell proliferation. Briefly, 5 * 10^4^ UMSCs per well were inoculated onto 14 mm glass slides and incubated overnight, and DNA synthesis was detected 48 hours after transfection. Nuclei were labeled with DAPI (Beyotime, China) under the guidance of the manufacturer’s instructions. and the proliferation of the cells was evaluated with a confocal laser scanning microscope (OLYMPUS, Japan). Statistical analysis was performed using ImageJ software.

### TUNEL assay

We used the TUNEL Apoptosis Detection Kit (Yeasen, China) to detect apoptosis under the guidance of the manufacturer’s instructions. 4 * 10^4^ UMSCs per well were inoculated onto 14 mm glass slides and incubated overnight, and apoptosis was measured 48 hours after transfection. Nuclei were labeled with DAPI, and apoptosis was evaluated with confocal laser scanning microscope (OLYMPUS, Japan). Statistical analysis was performed using ImageJ software.

### RT-PCR and qRT-PCR

We extracted total RNA with Ultrapure RNA Kit (CWBIO, China), and 1 μg RNA was used to synthesize cDNA with HiScript II Q RT SuperMix for qPCR (+gDNA wiper) (Vazyme, China). QRT-PCR was performed with Hieff® qPCR SYBR Green Master Mix (Yeasen, China) using CFX96 system (Bio-Rad, USA). The qRT-PCR primer sequences were available in Table 1. GAPDH was detected to homogenize samples.

**Table 1.**
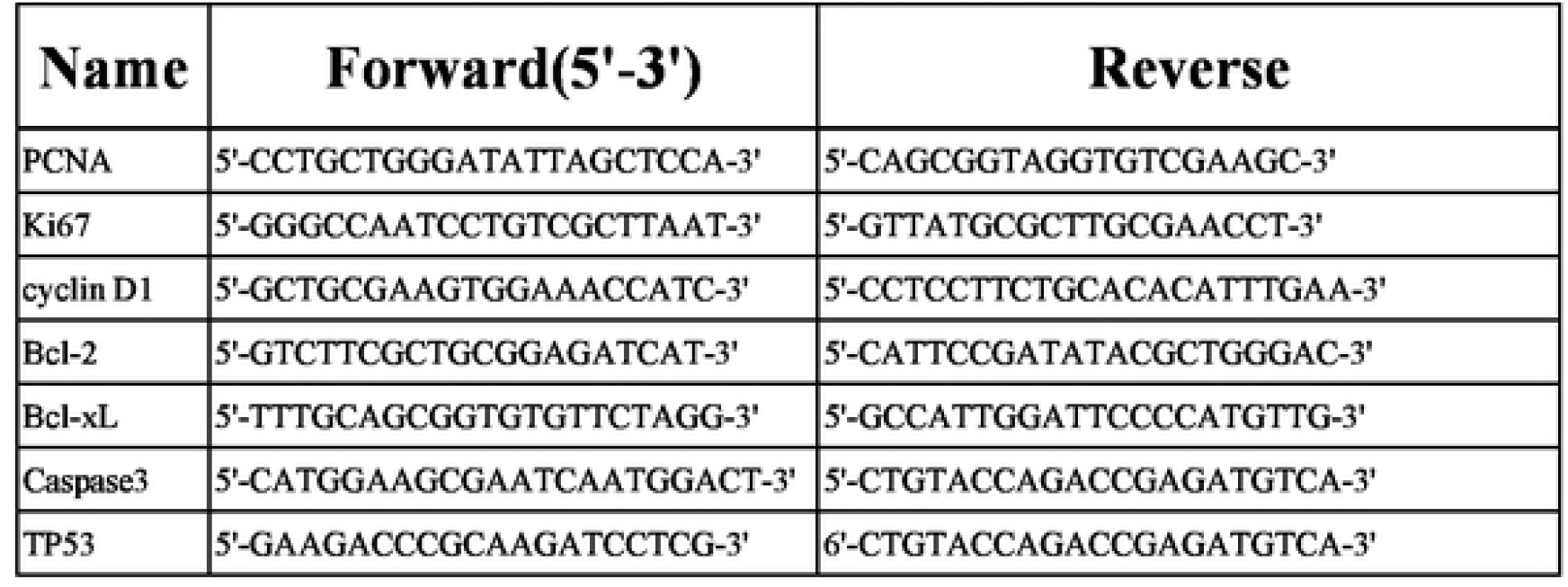
The primers used for qRT-PCR.

### WB assay

For WB, firstly lyses UMSCs with RIPA lysis buffer (Beyotime, China). Then separated 30 μg total protein according to the molecular weight by SDS-PAGE, then transferred to a PVDF membrane. After blocking the non-specific binding on the membrane with skim milk powder, incubated the membranes with the primary antibody for about 12 hours at 4 °C. All antibodies were as follows: Anti-GAPDH (sc-47724; Santacruz), Anti-CCND1 (ab68153; Abcam), Anti-Bcl-xl (2764; CST), Anti-Bcl2 (2872; CST), Anti-caspase-3 (ab13585; Abcam), Anti-PCNA (ab29; Abcam), Anti-Ki67 (ab197234; Abcam), Anti-p53 (ab1101; Abcam). The membrane was then incubated with a secondary antibody conjugated with HRP (Thermofisher, USA) for 1 hour at room temperature. Finally, the substrate (Thermofisher, USA) of the conjugated HRP was applied to the surface of the PVDF membrane and detected the chemiluminescence signal. GAPDH was detected to homogenize the sample.

### Luciferase assay

Luciferase assays were performed used a Bio-Glo™ Luciferase Assay System (Promega, USA). according to guidance of the manufacturer’s instructions. 5 * 10^4^ UMSCs per well were seeded in 24-well plates to detectthe luciferase activity 48 hours after transfection. Then measured the protein content using an Enhanced BCA Protein Assay Kit (Beyotime, China). The relative luciferase activity was characterized using the fluorescence value divided by the protein content.

### ChIP assay

We used Simple ChIP® Enzymatic Chromatin IP Kit (CST) to measure the binding of the ESR1 and TP53 promoters under the guidance of the manufacturer’s instructions. The DNA-protein complex was incubated with an anti-flag antibody (proteintech). The nucleic acid immunoprecipitated with the flag antibody was recovered, and the TP53 promoter fragment was detected by real-time PCR. The primers used were as follows: forward 5’-CCAGTCCTTACCATTTCTGC-3’, reverse 5’-GAGGCAGGCTCAGCCCTAGC-3’.

### Statistical analysis

Each experiment was repeated at least three times independently, and the data was expressed as mean ± SEM. Data were statistically analyzed by two-way ANOVA or by t-test. The significance level was characterized by the probability *P* < 0.05 and *P* < 0.01. * indicates *P* < 0.05, and ** indicates *P* < 0.01.

## Results

### ESR1 promoted the proliferation of UMSC

To explore whether ESR1 affects the proliferation of uterine fibroids, we overexpressed or knock down ESR1 in uterine smooth muscle UMSCs. We constructed a plasmid vector capable of overexpressing ESR1 and transfected into USMCs, transfected with pCDNA3.1 vectorplasmidwas a control group. The CCK-8 assays showed the cell viability of the UMSC cells was significantly promoted with the ectopic expression of ESR1 (Fig. 1A), the viability of the UMSC cells was decreased with the knockdown of endogenous ESR1 by siRNA (Fig. 1B). EdU-labeling assays indicated that ectopic expression of ESR1 accelerated division of USMCs (Fig. 1C), while ESR1 knockdown decreased the proliferation of USMC (Fig. 1D). Detecting the expression of proliferation marker genes with qRT-PCR and WB, data indicated that the proliferating marker genes PCNA, Ki67 and CCND1 was up-regulated after overexpression ESR1 (Fig. 1E and 1G), while the expression level of PCNA, Ki67 and CCND1 decreased with the knockdown of ESR1 (Fig. 1F and 1H).

**Fig.1.**
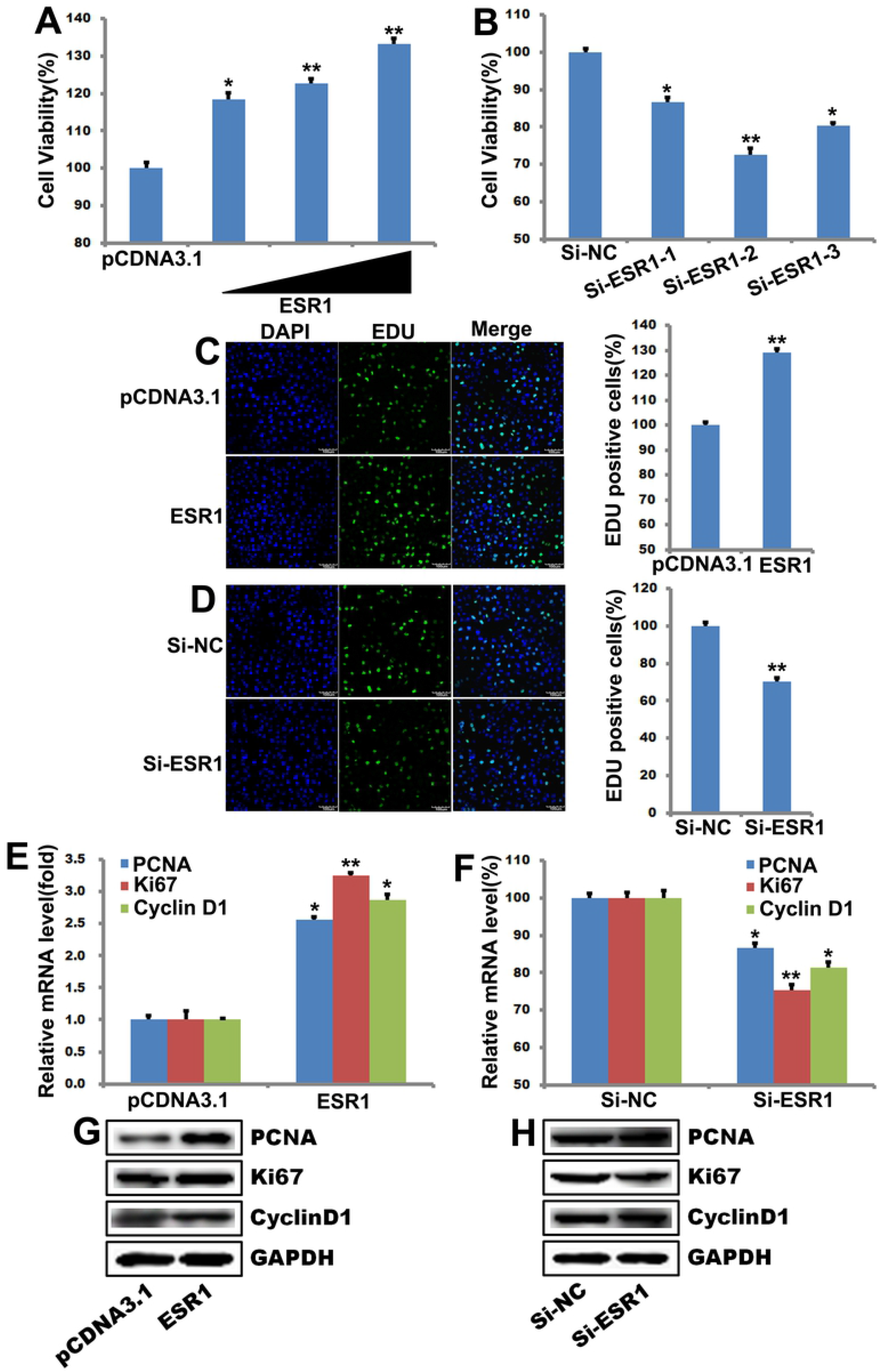
ESR1 promoted the proliferation of UMSC. (**A-B**) Cell viabilities of UMSC cells were detected by CCK-8 assay after transfected with 20, 40 or 60ng of ESR1 plasmid per well or pCDNA3.1, Si-ESR1 or Si-NC for 48 hours. The data represent mean ± SEM (n=6). (**C-D**) Cell proliferation of UMSC cells were detected by EdU assays after transfected with different 20, 40 or 60ng of ESR1 plasmid per well or pCDNA3.1, Si-ESR1 or Si-NC for 48 hours. (**E-F**) the mRNA levels in UMSC cells were detectedafter transfected with 20, 40 or 60 ng of ESR1 plasmid per well or pCDNA3.1, Si-ESR1 or Si-NC for 48 hours, and GAPDH was detected to homogenize samples. (**G-H**) The protein level in UMSC cells was tested after transfected with 20, 40 or 60ng of ESR1 plasmid per well or pCDNA3.1, Si-ESR1 or Si-NC for 48 hours, and GAPDH was detected to homogenize samples.

### ESR1 inhibited apoptosis in UMSC

We also explored the effect of ESR1 on the apoptosis of USMC by TUNEI. The results were showed in Fig. 2A and 2B, the result showed that overexpression of ESR1 remarkably inhibited the apoptosis compared to the control group, and knockdown of endogenous ESR1 increased the apoptosis of USMCs. We checked the level of Bcl-2, Bcl-xL and Cleaved-Caspase-3. The results showed that ectopic expression of ESR1 decreased the Cleaved-Caspase-3 and increased Bcl-2, Bcl-xL (Fig. 2C and 2E), and knockdown of endogenous ESR1 increased the Cleaved-Caspase-3 and decreased Bcl-2, Bcl-xL (Fig. 2D and 2F). In summary, all results indicated that ESR1 inhibited apoptosis in UMSCs.

**Fig. 2.**
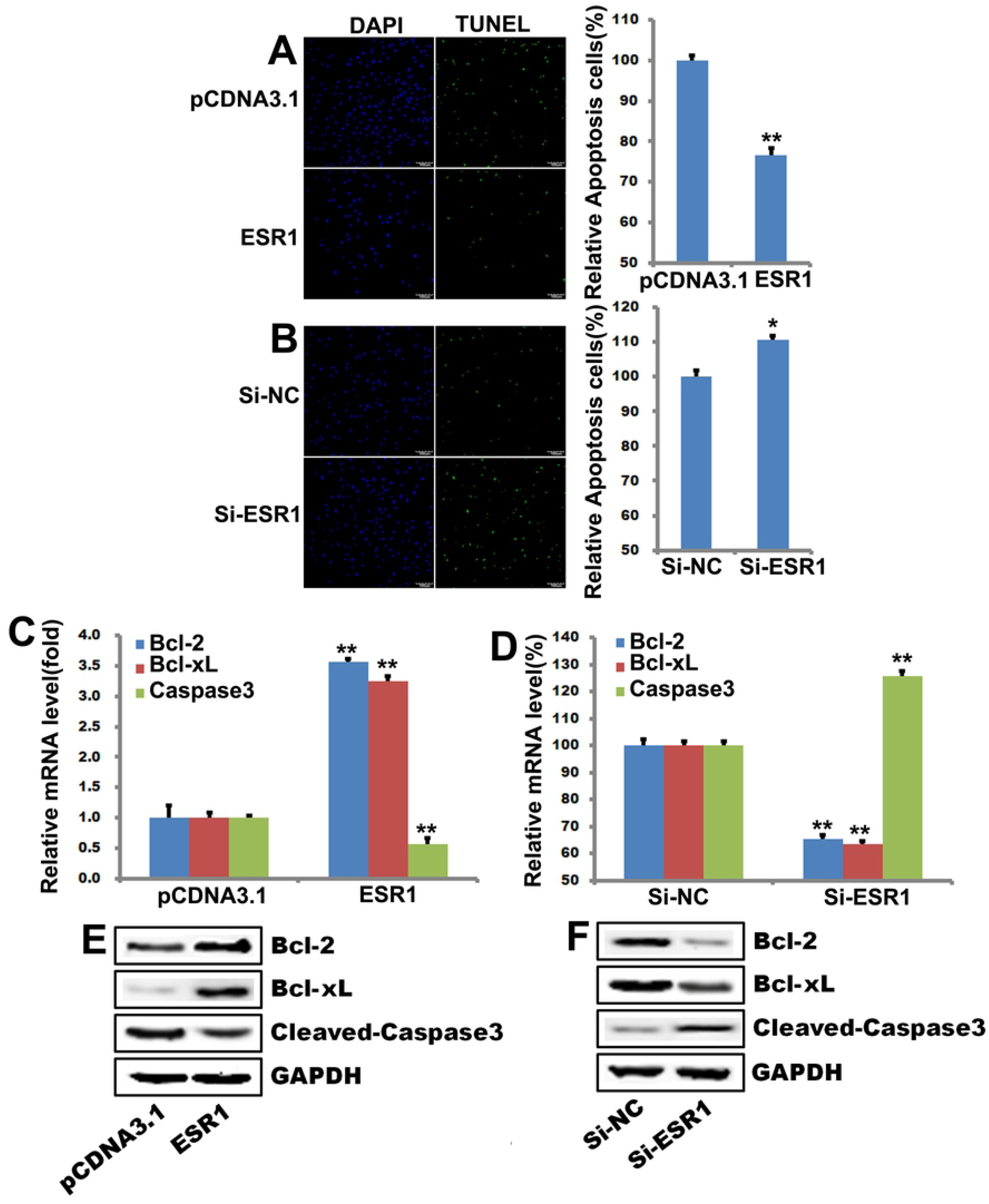
ESR1 inhibited apoptosis in UMSC. (**A-B**) Cell apoptosis of UMSC cells were detectedby TUNEL assays after transfected with ESR1 plasmid per well or pCDNA3.1, Si-ESR1 or Si-NC for 48 hours. (**C-D**) The mRNA levels in UMSC cells were detectedafter transfected with 20, 40 or60ng of ESR1 plasmid per well or pCDNA3.1, Si-ESR1 or Si-NC for 48 hours, and GAPDH was detected to homogenize samples. (**E-F**) The protein level in UMSC cells was detectedafter transfected with 20, 40 or 60ng of ESR1 plasmid per well or pCDNA3.1, Si-ESR1 or Si-NC for 48 hours, and GAPDH was detected to homogenize samples. The data represent mean ± SEM (n=3).

To address how ESR1 affect apoptosis, we first checked the expression of TP53 after ectopic expression or knockdown of ESR1 in USMCs. Our result showed that ectopic expression of ESR1 down-regulated TP53 and knockdown of ESR1 up-regulated TP53 (Fig. S1A-1D). To further confirm the mechanism of the inhibition effect of ESR1 on TP53, we checked ESR1 binging region inTP53 promoter. Bioinformatics analysis unveiled an ESR1 site on TP53 promoter. The wild-type TP53 promoter (TP53-WT-luc) and the MUT TP53 promoter (containing the mutant ESR1 binding sites) (TP53-M-luc) were cloned into pGL3 vector (Fig. S1E). Either the luciferase vectors TP53-WT-luc or the TP53-M-luc were co-transfected with ESR1 expression plasmid or empty vector pCDNA3.1 into UMSCs. We found that the ESR1 could inhibit the luciferase activity of TP53 WT luciferase vector while didn’t affect the activity of MUT vector (Fig. S1F). CHIP analysis also showed that ESR1 can bind directly to the TP53 promoter region (Fig. S1G). Together, these results confirmed that ESR1 inhibit TP53 expression by targeting the promoter.

### MiR-17-5p inhibited UMSCs’ proliferation

To explore whether MiR-17-5p affects the proliferation of uterine fibroids, miR-17-5p-mimics and miR-17-5p-inhibitor were transfected into USMCs. CCK-8 assay revealed that the cell viability of UMSCs was down-regulated as the transfection number of miR-17-5p mimics increased (Fig. 3A), the viability of the UMSCs was promoted after transfection miR-17-5p-inhibitor (Fig. 3B). EdU-labeling assays indicated that transfected of miR-17-5p-mimics inhibited USMCs’ proliferation (Fig. 3C), and inhibition of endogenous miR-17-5p promoted USMCs’ proliferation (Fig. 3D). Detection of expression of proliferation marker genes showed PCNA, Ki67 and CCND1 were down-regulated with the transfection of miR-17-5p-mimics in both mRNA and protein level (Fig. 3E and 3G), while the expression level of PCNA, Ki67 and CCND1 up-regulated with the inhibition of endogenous miR-17-5p (Fig. 3F and 3H).

**Fig. 3.**
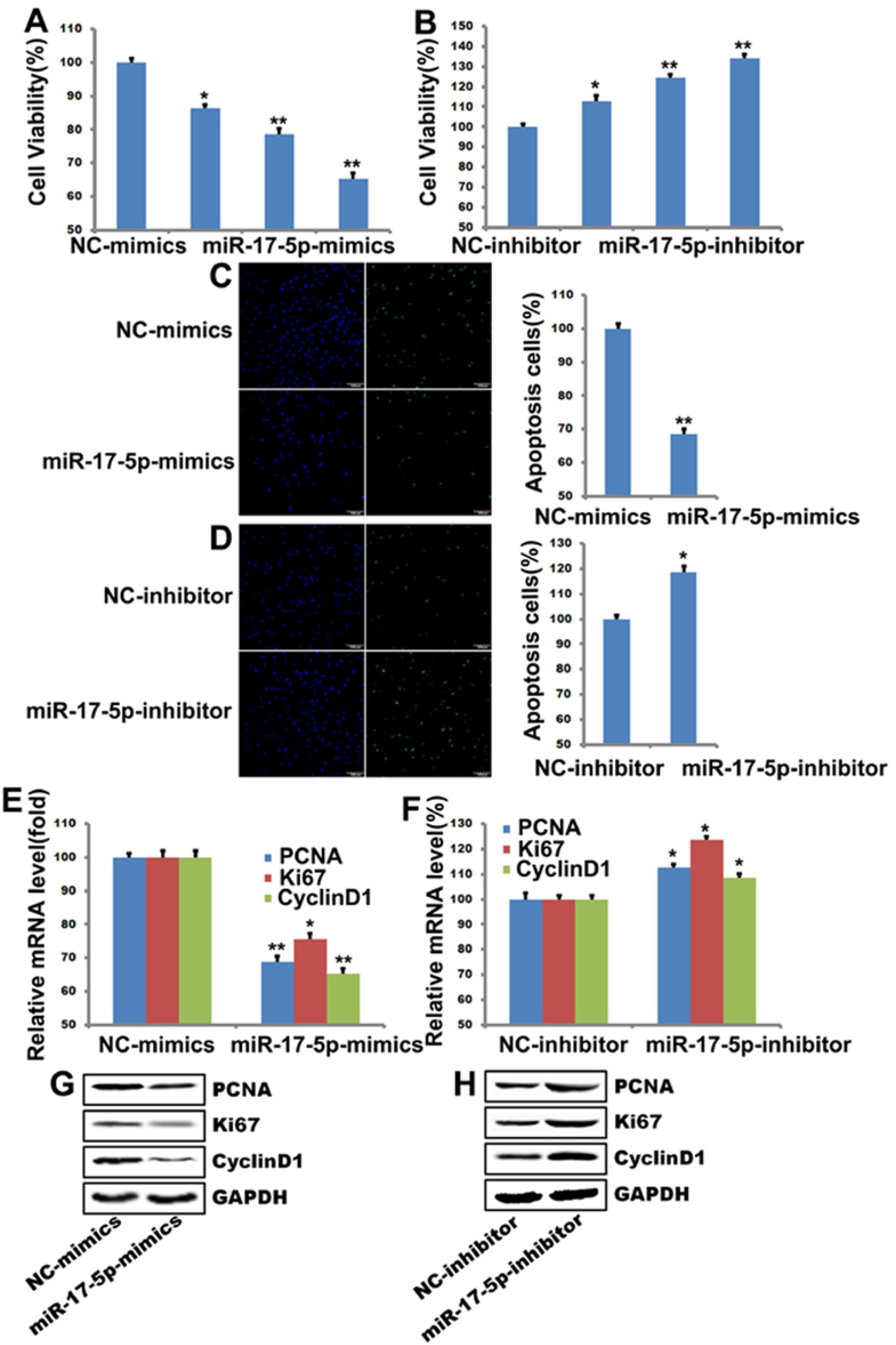
MiR-17-5p inhibited the proliferation of UMSC. (**A-B**) Cell viabilities of UMSC cells were detectedby MTT assays after transfected with miR-17-5p-mimics or NC-mimics, miR-17-5p-inhibitor or negative NC-inhibitor for 48 hours. (**C-D**) Cell proliferation of UMSC cells were detected by EdU assays after transfected with miR-17-5p-mimics or NC-mimics, miR-17-5p-inhibitor or negative NC-inhibitor for 48 hours. (**E-F**) The mRNA levels in UMSC cells were detectedafter transfected with miR-17-5p-mimics or NC-mimics, miR-17-5p-inhibitor or negative NC-inhibitor for 48 hours, and GAPDH was detected to homogenize samples. (**G-H**) The protein level in UMSC cells was detectedby WB after transfected with miR-17-5p-mimics or NC-mimics, miR-17-5p-inhibitor or negative NC-inhibitor for 48 hours, and GAPDH was detected to homogenize samples.

### MiR-17-5p promoted USMCs’ apoptosis

We also explored whether miR-17-5p affect USMCs’ apoptosis via TUNEI. The data indicated that transfection of miR-17-5p remarkably promoted the apoptosis progression, and inhibition of endogenous miR-17-5p decreased the apoptosis of USMCs. We checked the expression of Bcl-2, Bcl-xL and Cleaved-Caspase-3. The results showed that transfection of miR-17-5p increased the Cleaved-Caspase-3 and decreased Bcl-2, Bcl-xL (Fig. 4C and 4E), and inhibition of endogenous miR-17-5p decreased the Cleaved-Caspase-3 and increased the expression of Bcl-2, Bcl-xL (Fig. 4D and 4F). All results indicated that miR-17-5p promoted apoptosis in UMSCs.

**Fig. 4.**
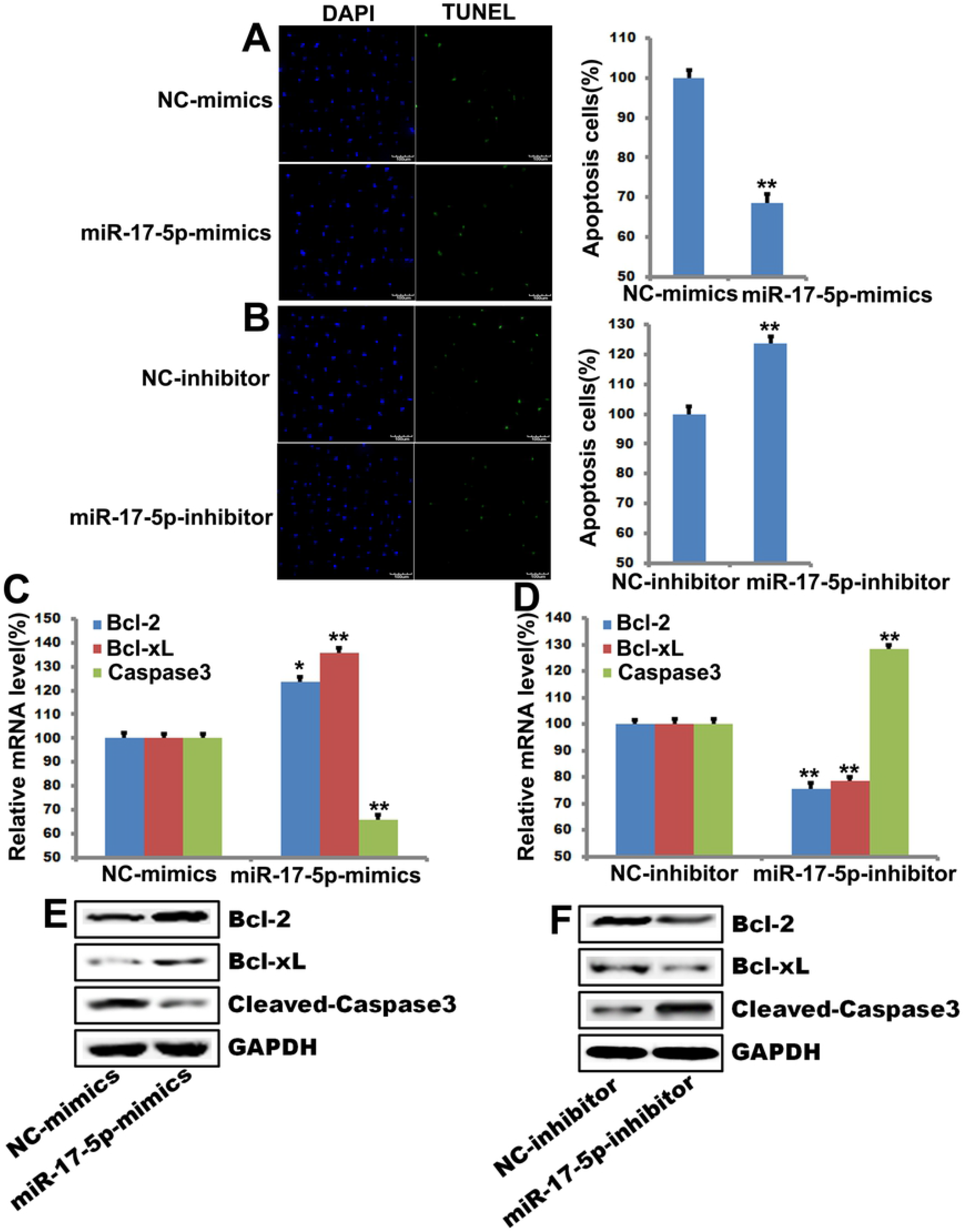
MiR-17-5p promoted apoptosis in UMSC. (**A-B**) Cell apoptosis of UMSC cells were detectedby TUNEL assays after transfected with miR-17-5p-mimics or NC-mimics, miR-17-5p-inhibitor or negative NC-inhibitor. (**C-D**) The mRNA levels in UMSC cells were detectedafter transfected with miR-17-5p-mimics or NC-mimics, miR-17-5p-inhibitor or negative NC-inhibitor, and GAPDH was detected to homogenize samples. (**E-F**) The protein level in UMSC cells wasdetected after transfected with miR-17-5p-mimics or NC-mimics, miR-17-5p-inhibitor or negative NC-inhibitor for 48 hours, and GAPDH was detected to homogenize samples.

### MiR-17-5p antagonizes the role of ESR1 in promoting proliferation and inhibiting apoptosis

To explore the relationship between ESR1 and miR-17-5p on the effect of proliferation and apoptosis in USMCs, we transfected miR-17-5p mimics while overexpressing ESR1. Cell proliferation was detected by CCK-8 assays and EdU-labeling assays, and apoptosis was detected by TUNEI. The results showed that the transfection of miR-17-5p mimic could partially inhibit ESR1-promoted USMCs ability and proliferation (Fig. 5A and 5B). Similarly, the transfection of miR-17-5p mimic could also antagonize ESR1-mediated USMCs apoptosis (Fig. 5C). We checked the expression of PCNA, Ki67, CCND1, Bcl-2, Bcl-xL and Cleaved-Caspase-3. Our data showed that the transfection of miR-17-5p mimic could also antagonize ESR1-mediated expression of proliferation and apoptosis relative gene (Fig. 5D and 5E).

**Fig. 5.**
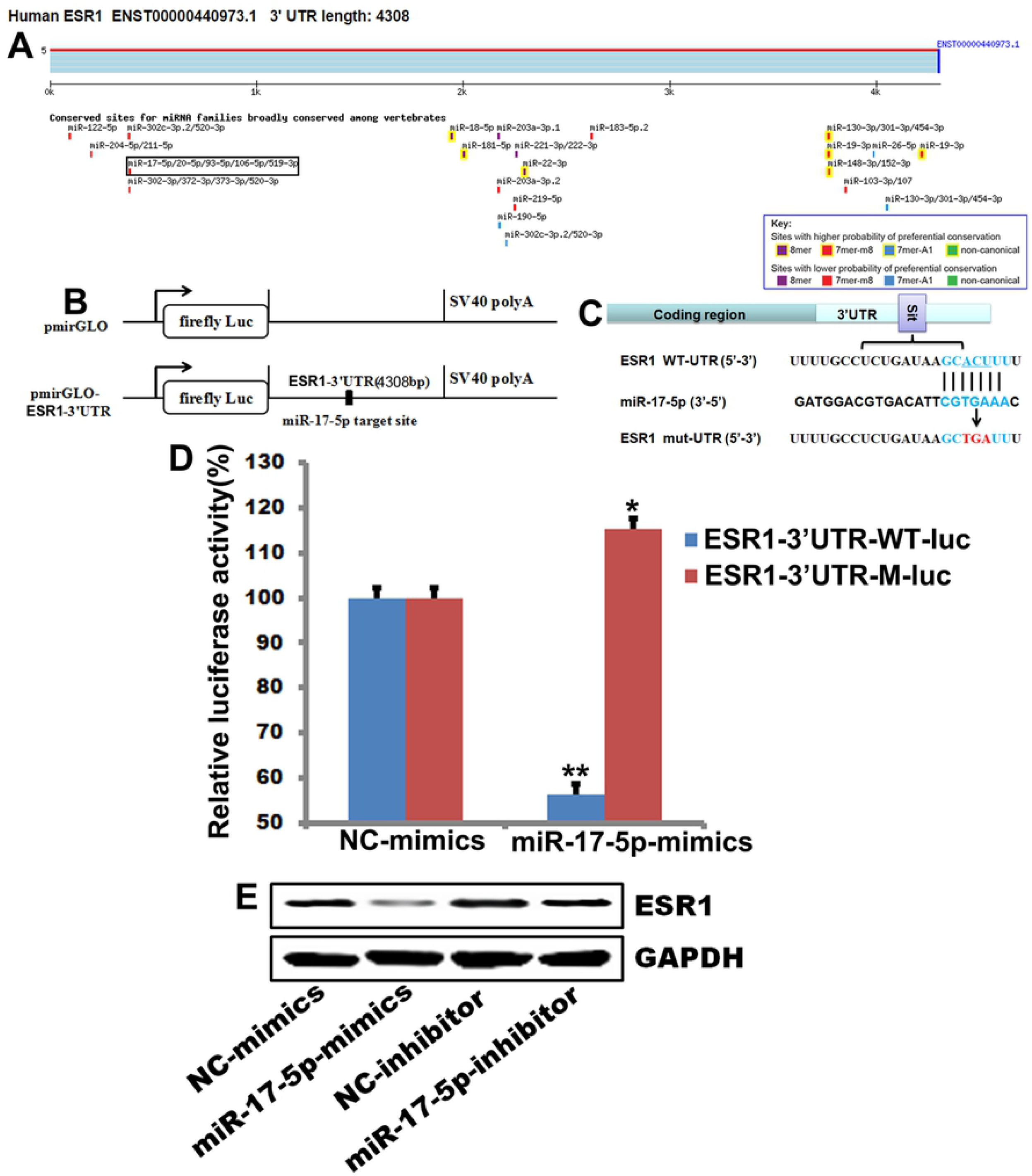
MiR-17-5p antagonizes the role of ESR1 in promoting proliferation and inhibiting apoptosis. Co-transfecting with miR-17-5p-mimics or NC-mimics and Si-ESR1 or Si-NC to UMSC cells for 48 hours. (**A**) Testing the cell viabilities of with MTT assays after transfection. (**B**) Cell proliferation of UMSC cells were detected with EdU assay. (**C**) Cell apoptosis of UMSC cells were detectedby TUNEL assays after transfection. (**D**) Testing mRNA level after transfection, and GAPDH was detected to homogenize samples. (**E**) The protein level in UMSC cells was detectedafter transfection, and GAPDH is detected to homogenize samples.

To explore how miR-17-5p affects the expression of ESR1, we used targetscan website to predict the 3’UTR of ESR1. The analysis showed there was a miR-17-5p specific site (Fig. S2A). To explore whether miR-17-5p binds to ESR1’s 3’UTR, two luciferase vectors containing either the WT ESR1 3’UTR (ESR1-3’UTR-WT-luc) or the MUT ESR1 3’UTR (ESR1-3’UTR-M-luc) (Fig. S2B and S2C) were constructed. Then co-transfected ESR1-3’UTR-WT-luc or ESR1-3’UTR-M-luc with miR-17-5p-mimic or NC-mimic. We found that the miR-17-5p mimic could down-regulate the activity of the ESR1 WT luciferase vector while didn’t affect the activity of MUT vector (Fig. S2D). And miR-17-5p mimic decreased the expression of ESR1, and miR-17-5p inhibitor increasedthe expression of ESR1 (Fig. S2E). Together, allresults showedthat miR-17-5p inhibited ESR1 expression in UMSCs by targeting ESR1 3’UTR.

## Discussion

Uterine fibroids are benign lesions or uterine tumors.[1] At present, the main treatment of uterine fibroids is hysterectomy, which may affect fertility and have a profound impact on health.[2] The development of new uterine fibroids treatments and drugs depends on a deeper understanding of the development of uterine fibroids and the discovery of new therapeutic targets.

Tumor cells were uncontrolled proliferative state.[11] Previous research showed that ESR1 was linked to proliferation of breast cancer.[12] Here, our data showed that expression of the proliferating marker genes PCNA, Ki67 and CCND1 was increased in the overexpressed ESR1 group in USMC cells (Fig 1G).

We also reported that ESR1 inhibited apoptosis in UMSC (Fig. 2A), ectopic expression of ESR1 decreased the Cleaved-Caspase-3 and up-regulated Bcl-2, Bcl-xL (Fig. 2C and 2E), and knockdown of endogenous ESR1 increased the Cleaved-Caspase-3 and decreased Bcl-xL, Bcl-2 (Fig. 2D and 2F). In addition, ESR1 bound to promoter of TP53 and inhibit TP53 expression (Fig. S2).

MiR-17-5p was reported to have important role in breast cancer.[7,8,6,13] Here, we reported that miR-17-5p had an effect of inhibiting proliferation and promoting apoptosis in USMCs (Fig. 3 and Fig. 4).

MiRNAs negative regulation of target genes via binding to mRNA or 3’UTR to inhibit the expression of target genes.[5,14,15] Here, miR-17-5p was proved to target 3’UTR of ESR1 to inhibit its expression and antagonize its function (Fig. S1).

## Acknowledgements

This work was supported by Projects of medical and health technology development program in Shan Dong Province (2017WS269) and Academy-level Science and Technology Program of Shandong Academy of Medical Sciences (2019-19).

## Author contributions

Kun Li designed research; Lan Luo and Jian-Jun Cao performed research; Hui-Min Zhang, Kun Chen and Lan Luo analyzed data; Lan Luo, Xing-Hua Liao and Kun Li wrote the paper.

---

| **Abbreviation** | **Full name** |
| --- | --- |
| ESR1 | Estrogen receptor 1 |
| miRNAs | microRNAs |
| 3'UTR | 3' untranslated region |
| mRNA | messenger RNA |
| USMCs | Uterine smooth muscle cells |
| ATCC | American Type Culture Collection |
| DMEM | Dulbecco's modified Eagle's medium |
| FBS | fetal bovine serum |
| siRNAs | small interfere RNAs |
| NC | negative control |
| cDNA | complementary DNA |
| WT | wild type |
| MUT | mutant |
| CCK-8 | Cell Counting Kit-8 |
| EdU | 5-ethynyl-2'-deoxyuridine |
| DAPI | 4,6'-diamidino-2-phenylindole |
| TUNEL | Terminal-deoxynucleotidyl transferase mediated nick end labeling |
| RT-PCR | Reverse-transcription polymerase chain reaction |
| qRT-PCR | quantitative real-time RT-PCR |
| WB | Western blot |
| RIPA | Radio Immunoprecipitation Assay |
| SDS-PAGE | sodium dodecyl sulfate-polyacrylamide gel electrophoresis |
| PVDF | polyvinylidene fluoride |
| HRP | horseradish peroxidase |
| ChIP | Chromatin immunoprecipitation |
| SEM | standard error of the mean |
| ANOVA | analysis of variance |

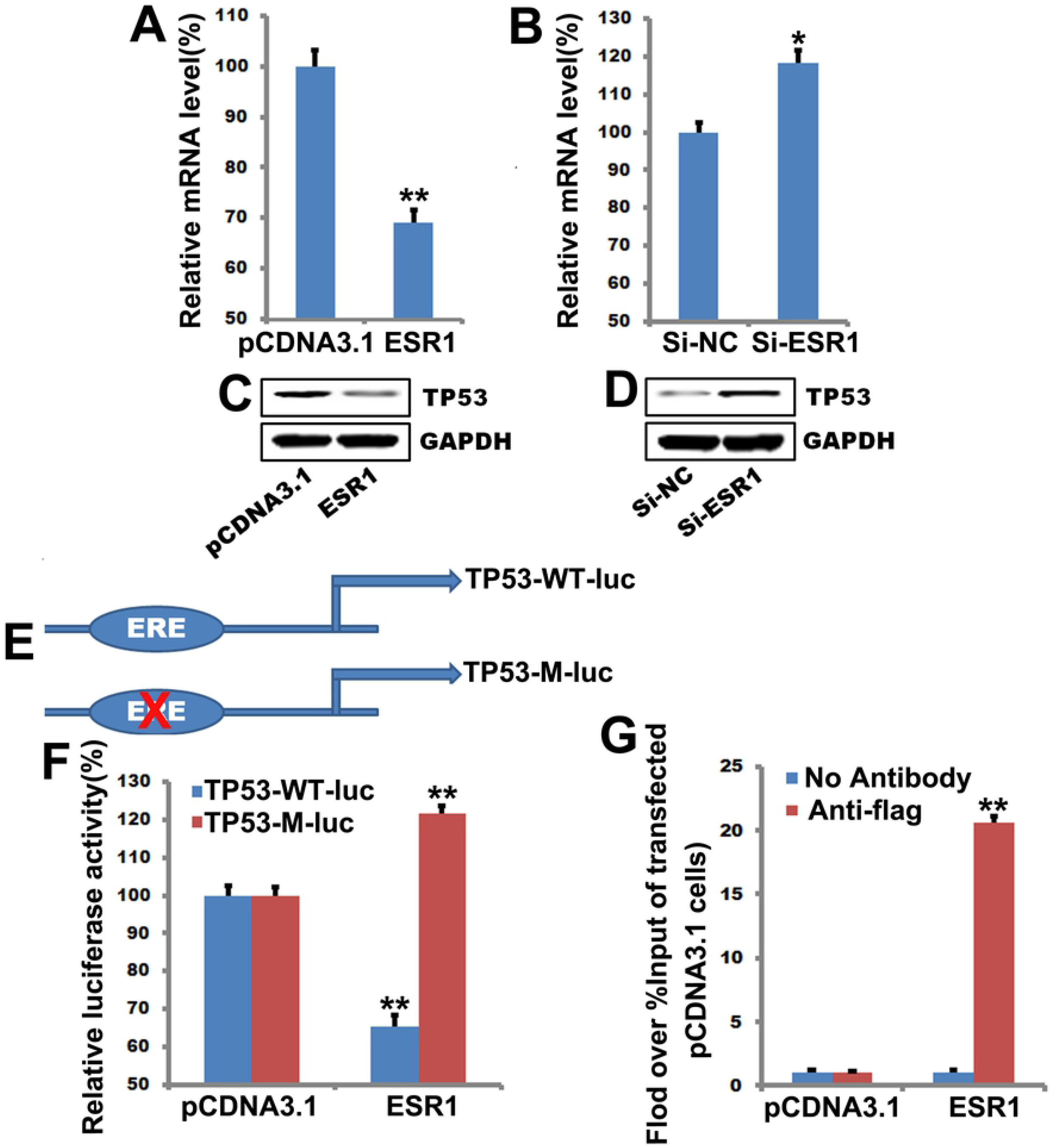

